# Fully Interpretable Deep Learning Model of Transcriptional Control

**DOI:** 10.1101/655639

**Authors:** Yi Liu, Kenneth Barr, John Reinitz

## Abstract

The universal expressibility assumption of Deep Neural Networks (DNNs) is the key motivation behind recent work in the system biology community to employ DNNs to solve important problems in functional genomics and molecular genetics. Because of the black box nature of DNNs, such assumptions, while useful in practice, are unsatisfactory for scientific analysis. In this paper, we give an example of a DNN in which every layer is interpretable. Moreover, this DNN is biologically validated and predictive. We derive our DNN from a systems biology model that was not previously recognized as having a DNN structure. This DNN is concerned with a key unsolved biological problem, which is to understand the DNA regulatory code which controls how genes in multicellular organisms are turned on and off. Although we apply our DNN to data from the early embryo of the fruit fly *Drosophila,* this system serves as a testbed for analysis of much larger data sets obtained by systems biology studies on a genomic scale.

## 1 Introduction

A central unsolved problem in biology is to understand how DNA specifies how genes turn on and off in multicellular organisms. The “universal expressibility” of deep neural nets suggests that they might be a valuable tool in this undertaking, but their applicability and acceptance in solving problems in natural science has been limited by the uninterpretability of their internal computations. We address both of these areas in this report by describing the reimplementation of a specific and highly predictive model of transcriptional control as a deep neural net (DNN). The model, chemical in nature, has a feedforward mathematical structure in which every layer has a specific scientific interpretation. When translated into DNN formalism, it provides an example of a DNN in which the internal structure is well understood and which may be of value to the machine learning field.

Deep learning has been widely deployed in genomics and systems biology over the last few years [11, 2, 27, 37, 9, 43, 39, 13, 35, 34]. Many of the developed tools have been highly successful in classification problems such as the identification of binding sites, open regions of chromatin, and the location of enhancers. Deeper understanding requires more quantitative studies. One recent example that goes beyond classification concerns a fully quantitative and highly predictive DNN model of the role of untranslated RNA leader sequences in gene expression in yeast [11], We believe that these studies have two sets of limitations. First, they take a universal expressibility approach without much understanding and interpretation of the underlying chemical and biological mechanisms giving rise to to the phenomena under study. This limits the contributions DNNs can make to increasing human understanding of fundamental biological processes. Here we consider a DNN in which each layer has a specific chemical or biological interpretation. Studies of regulatory DNA with DNNs have treated only the sequence itself, but in metazoa (multicellular animals) different cell types have very different gene expression states although they contain the same DNA. Here we consider a DNN in which state is described not only by the sequence, but also by the set of regulatory proteins present in each cell. We now describe the problem to be solved in both mathematical and biological terms.

Mathematically, the “expression” of a gene is the rate at which it synthesizes mRNA (the “transcript”) from a complementary DNA template. This rate *d*[mRNA]/*dt* = *f* (***D**, v*) where ***D*** = (*D_i_*), where each *D_i_* ∈ {*A, C, G, T*} is a base in the sequence of regulatory DNA, and *v* = (*v*_1_,…, *v_a_*,…, *v_n_*), where each *v_a_* is the nuclear concentration of a regulatory DNA binding protein known as a “transcription factor” (TF). The machine learning task is to learn the function *f* from a series of observations (***D**_j_*, *v_jk_*) of the expression of sequence *j* in cell type *k*, where *k* ∈ {1,…, *M*}. The essence of the scientific problem is that each sequence ***D**_j_* must express correctly in each cell type, reflecting the fact that in a multicellular organism, different sets of genes are expressed in different cell types, but each cell type contains the same DNA.

Biologically, regulatory DNA is noncoding DNA which can be upstream (5’), downstream (3’), or within (intronic) the complementary mRNA template, but it is distinct from (exonic) DNA that contains codons that specify the amino acid sequence of the gene’s protein product. TFs bind to DNA in its double stranded form, in which each strand has a complementary base (A and T; G and C) at each position and the two strands have opposite 3’-5’ orientations. In metazoans, regulatory DNA is frequently much larger than the coding portion of the gene. The regulatory DNA contains segments of 500 to 1000 base pairs (bp) called “enhancers”, each of which directs expression in a particular domain or tissue type. In this study, we consider a gene called *eve* which acts in the early embryo of the fruit fly *Drosophila melanogaster*, at which time it forms a pattern of seven stripes as shown in Figure 1. The entire gene is 16.5 kilobases (kb) of DNA in length, but the mRNA transcript is only 1.5 kb (Figure 2). We consider the action of *eve* from 1 to 3 hours after the start of embryonic development. At this time the embryo is a hollow ellipsoid of cell nuclei that can be treated like a naturally grown gene chip in which d[mRNA]/*dt*, ****D**_j_**, and *v_jk_* are fully observable at a quantitative level. The embryo contains two orthogonal axes in the anterior-poster (A-P) and dorsal-ventral (D-V) directions. In the central portion of the embryo, gene expression on the two axes is uncoupled, so cell type and gene expression can be visualized by plotting relevant state variables in one dimension.

**Figure 1:**
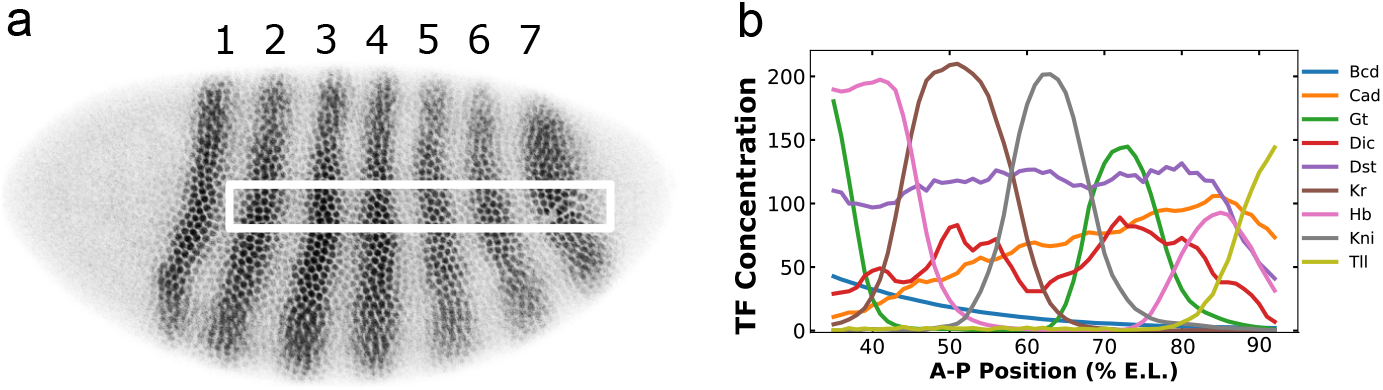
(a) shows a *Drosophila* embryo about 3 hours after fertilization which has been stained for Eve protein as described [48]. Anterior is to the left and dorsal is up. The dark shades indicate the concentration of Eve protein and the stripes are numbered. The white box is the one dimensional region of interest used to generate the data[20]. (b) The TF concentrations found across the embryo [48]. In the graph, 58 data points are shown, corresponding to 58 nuclei on the A-P axis. Each nucleus is 1 % E.L. in size. The identity of TFs is shown in the key; the horizontal axis shows position in percent egg length (E.L.) and the vertical axis shows protein concentration.

In this paper, we show that DNNs can be used to generate a predictive model of gene expression. Our starting point is a previously published model of transcriptional control [23], which is one of a family of so-called thermodynamic transcription models [38, 21, 42, 40, 22, 16, 23, 32, 41, 4, 5]. In these models, occupancy of DNA by TFs is calculated using thermodynamics, and phenomenological rules are used to calculate the transcription rate from the configuration of bound factors. Like DNNs, thermodynamic transcription models have a feedforward structure that can be described in layers. The form of the resulting equations makes back propagation and hence SGD difficult or impossible because of the need to hand code complex partial derivatives, so these models are optimized by zero order methods such as Simulated Annealing or Genetic Algorithms. Despite this apparent mathematical distinction, we were able to translate the chemical model of transcription into a standard DNN form that could perform rapid learning by SGD.

**Figure 2:**
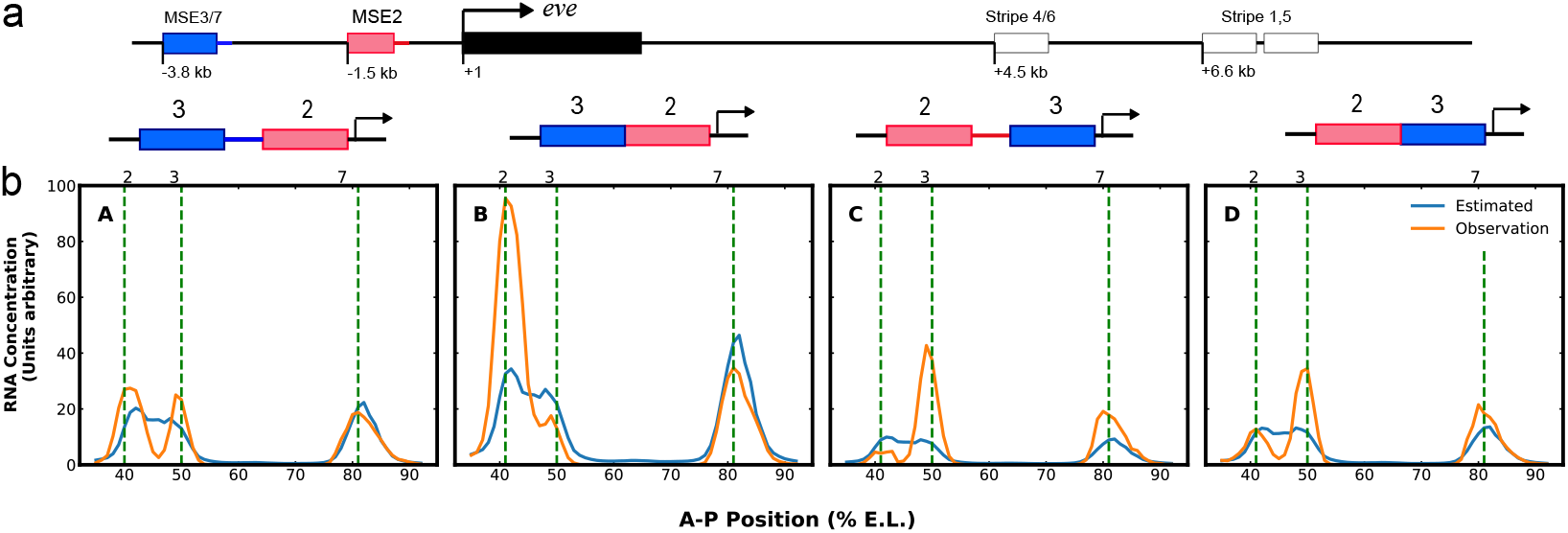
(a): The figure shows a diagram of the eve locus. The transcript is indicated by black box; enhancers are indicated by pink, blue or white boxes and are labeled with the eve stripe that they drive. (b): Fusion constructs that are used in the training process. The blue box represents the MSE3/7 enhancer and the red box represents the MSE2 enhancer. The four graphs in (b) shows the data used for training and outcome from the model after 500 epochs using Adam. In each graph, 58 data points are shown, corresponding to 58 nuclei on the A-P axis. Each nucleus is 1 % E.L. in size. The orange lines show the observed data from the experiments [23] and the blue lines show the model output. The overall RMS is of the training is calculated to be 10.4.

The resulting model is, to our knowledge, one of the first fully interpretable DNNs with an exact interpretation for each of the unknown parameters. It is also an example of a biologically validated DNN that is not perceptron based. Some of the parameters can be and are extracted from independent experiments, resulting in a very small number of parameters for an extremely deep network. Although theories of metazoan transcriptional control have largely been developed in the fruit fly *Drosophila* because of its unique experimental advantages, they have also been applied to mouse [6]. Below, in Section 2, we will describe the function of each layer and the interpretation of parameters. In section 3, we focus on the training and evaluation of performance. Finally, in section 4, we discuss the scientific implications of our results.

## 2 Understanding each layer

The model’s input is DNA sequence and TF concentration. The TFs have functional roles which, in the present application, are known from independent experiments. Activators activate transcription, quenchers suppress the action of activators over a limited range, and certain activators can also convert nearby quenchers into activators. In each of these regulatory mechanisms, multiple bound TFs are required to perform a regulatory action. We perform this calculation as follows. Binding site locations and affinities are determined from sequence. Together with TF concentrations, this information is used to calculate the occupancy of each binding site. We then calculate the effects of coactivation, followed by the effects of quenching. The total amount of remaining activation is then summed and passed through a thresholding function. We describe each step below.

### 2.1 Computing Fractional Occupancy

#### 2.1.1 Identifying the binding sites

An indicator representation of DNA sequence is used as input. The column index is the base pair position number in the sequence. The row index indicates which of the 4 bases (A,C,G,T) is observed. In the case where the sequence cannot be identified, a fifth row (N) is used. For example, if we have a sequence of ACTNGTTA, the corresponding matrix is

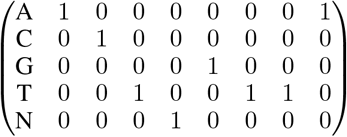

The nine TFs of interest are Bicoid (Bcd), Caudal (Cad), Drosophila-STAT (Dst), Dichaete (Dic), Hunchback (Hb), Krupple (Kr), Knirps (Kni), Giant (Gt), and Tailless (Tll) (Figure 1b). The identification and affinity characterization of binding sites for TF *a* requires a convolution layer with a Position Weight Matrix (PWM_*a*_) as its kernel. The PWM can be understood as a convolution kernel in which the number of columns is the number of nucleotides of DNA in physical contact with a bound TF. Padding is not required for the columns. In the work described here, we use the same high quality experimentally obtained PWMs as previously [23]. Unlike many convolution matrices used in deep learning, PWMs make direct experimental predictions about DNA properties, a fact used in a previous study to experimentally confirm PWMs obtained by deep learning techniques [11].

Chemically, the PWM represents an additive model of binding in which the Gibbs free energy Δ*G* of binding is the sum of the free energies of binding to each nucleotide. Statistically, the PWM score can be regarded as the likelihood of finding a binding site at a given position, calculated using a variational approximation of the likelihood using only the marginal likelihood of each base. The resulting score, denoted by *S*_*i,i+m;a*_, is an affine transformation of the the free energy △G_i,i+m;α_ of TF *a* binding to a site extending from base *i* to base *i* + *m*. In many cases, including most of the TFs considered here, m can be read off directly from DNAse I footprints [44, for example]. TFs physically bind in the major groove of the DNA double helix but the PWM is convolved with only a single strand. We compensate for this fact by also scoring the complementary strand, in which bases are replaced by their complements (A → T; C → G; T → A; G → C), and orientation is reversed. Scoring each strand results in a 1 × *n* array of scores S_α_, where n is the sequence length. At each base position, we set the score to be the larger of the two scores at that position.

We next calculate the equilibrium affinity *K_i,i+m;a_* of each binding site. If Δ*G_i,i+m;a_* were known in units of kcal/mole, then *K_i,i+m;a_* = exp(Δ*G_i,i+m;a_/RT*), where *R* is Boltzmann’s constant and *T* the absolute temperature. *S_i,i+m;a_* is related to Δ*G_i,i+m;a_* by an affine transformation *ax* + *b* in which *b* is found from experiment and a is learned by training as follows. Our convolution kernel is accompanied by *a* fixed bias *S_max;a_* which is the maximum possible score for any particular TF *a*. We only consider binding to sites with scores greater than zero, so we pass the exponentiated free energy through a `ReLU` and apply an indicator function to assure that below threshold sites disappear, so that

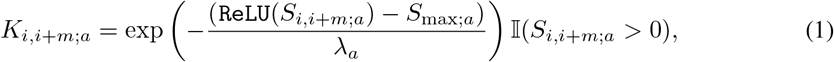

where λ_*a*_ is a learnable parameter for each TF *a*. The *K_i,i+m;a_*, arranged according to positions on the DNA sequence, produce another 1 × (*n - m*) vector *K_a_*. Adjusting for the size of *m*, we further concatenate *K_a_* for all TFs to produce *K*, which we use for the calculation of fractional occupancy.

#### 2.1.2 Belief propagation and partition function computation

Moving from affinity *K_i,i+m_* to fractional occupancy requires the consideration of all possible states in which the TFs can bind on the DNA. The fractional occupancy *f_i,i+m;a_* denotes the average occupancy of the site at equilibrium or the probability of finding the specific protein a at the site (*i, i + m*) at a given instant. The calculation requires consideration of interactions between sites. Two overlapping sites cannot be occupied at the same time, and in some cases a TF bound at one site increases the binding affinity at a nearby site by a factor of 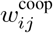. In the present application, cooperativity only occurs between pairs of bound Bcd less than 60 bp apart. This calculation is best performed by computing the partition function *Z*, which can be calculated by a fast, recently discovered algorithm [4, Appendix S1]. The algorithm is essentially a form of Belief propagation which can be represented as a bi-directional RNN across *K*, and is given in Algorithm 1.

##### Algorithm 1

The Algorithm for Fractional Occupancy

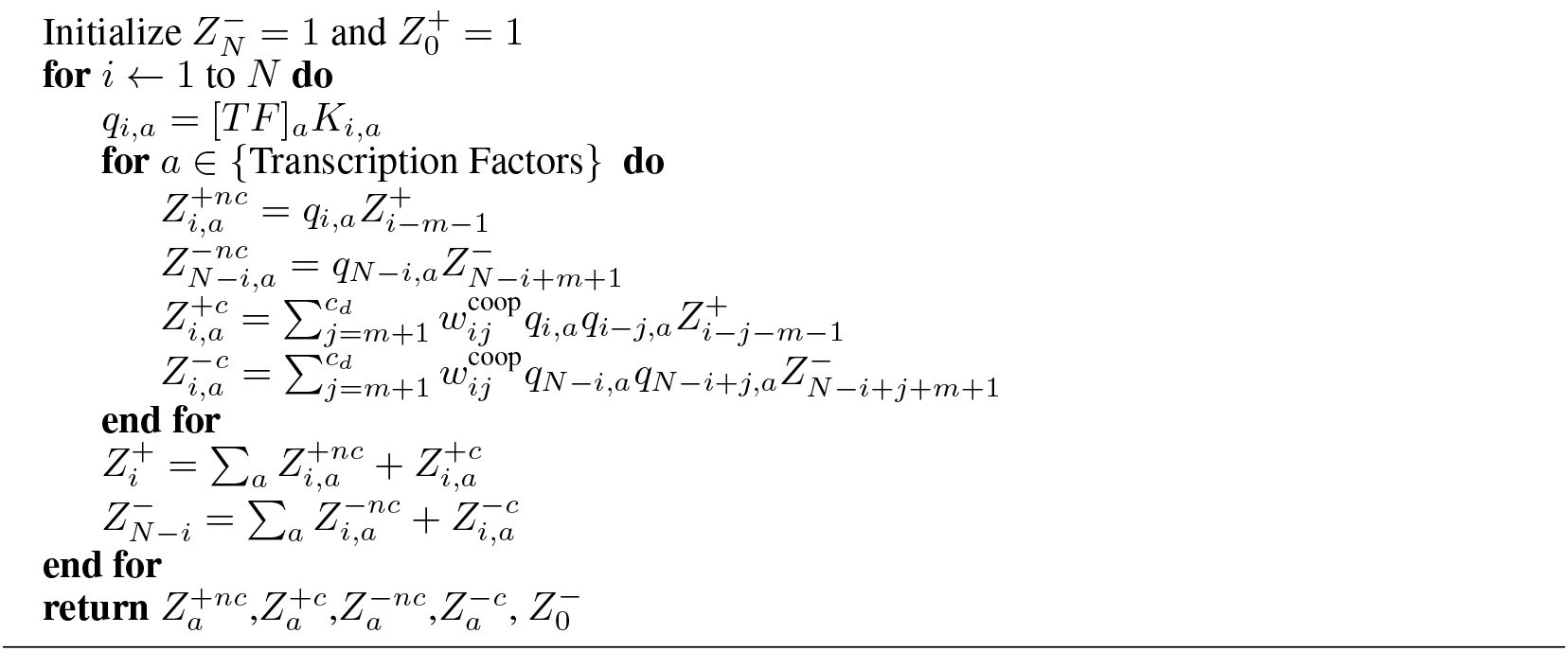

The fractional occupancy is then given by

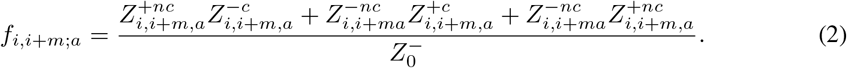

### 2.2 TF-TF Interactions

Interactions between bound TFs follow phenomenological rules. A central feature of the cis-regulatory DNA of metazoan genes is the fact that biological function is encoded in multiple binding sites [47, 44, 45]. This fact is expressed mathematically in the phenomenological equations below. In the present application, bound TFs have specific roles derived from specific experimental results, although this approach also works if the roles are not known a priori [6]. Here, Bcd, Cad, Dst and Dic are activators; Hb, Tll, Kni, Kr and Gt are quenchers; and both Bcd and Cad are coactivators of Hb. We now describe the actions of each class of TF in the order which we compute them. The ultimate goal of this computation is to obtain the summed action of activators after their contribution has been increased by coactivation and diminished by quenching.

#### 2.2.1 Coactivators

Coactivators turn a nearby quencher into an activator. This action is described by the equation [23, 4]

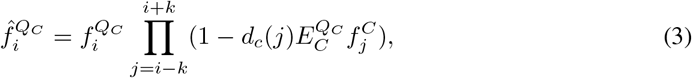

where 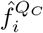 is the portion of activator fractional occupancy created from the total fractional occupancy 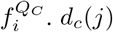 is a convolutional kernel describing the distance dependence of coactivation. *d_c_*(*j*) = 1 if |*j – i*| less than 156 bp. In this application, *k* = 206. As |*j – i*| increases to 206 bp, *d_c_*(*j*) decreases linearly to 0. 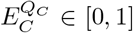 denotes the relative strength of the coactivators. 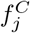 simply refers to fractional occupancy of Bcd and Cad since they are the only coactivators in this setting.

It is easy to observe that equation (3) is the first term of a Taylor expansion of

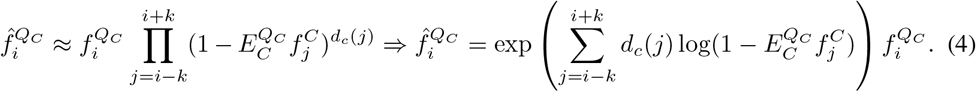

This can be turned into three convolutional linear activation units, so that

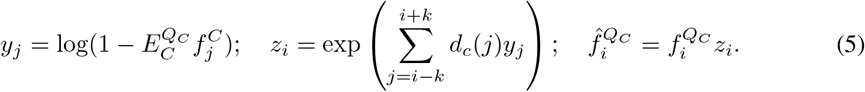

#### 2.2.2 Quenchers

As their name suggests, quenchers suppress activation. Their action is local and occurs only within around 100 to 150 base pairs [17]. The basic mathematical formulation of the effect of nearby quenchers on the activator at position *i* is given by [23]

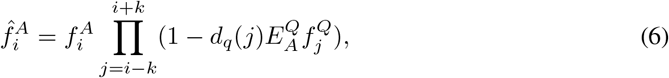

where 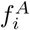 is the fractional occupancy of activators whether coactivated or not, 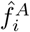 is the fractional occupancy of activators after quenching, 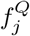 is the fractional occupancy of a quencher bound at position 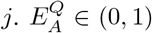 is the strength of quencher *Q* on activator *A* and *d_q_*(*j*) is a convolutional kernel representing the range of quenching on the DNA strand. *k* = 150 bp, and *d_q_* (*j*) = 1 when |*j – i*|≤ 100 and goes linearly down to 0 from 100 to 150 bp. Using a mathematical argument similar to that of the previous section, we can write

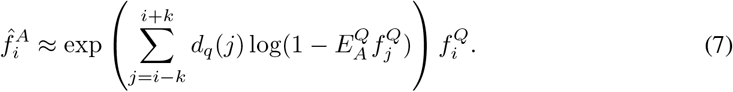

This gives three convolutional linear activation units.

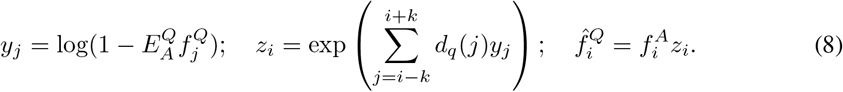

#### 2.2.3 Activation

Last, we sum the fractional occupancies of all activators remaining after the previous two steps. We consider the activators to lower the energy barrier in a diffusion limited Arrhenius rate law [4], which has the mathematical form of a sigmoidal thresholding function. This results an fully connected layer which yields the mRNA synthesis rate. In the experimental system used, mRNA has a life time much shorter than the time required to change transcription rates, so that the mRNA concentration [`mRNA`], an experimentally observable quantity, is given by

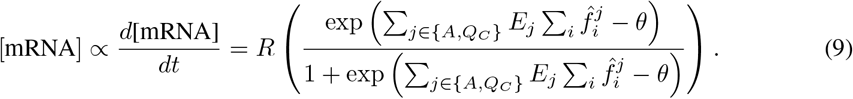

Here *E_j_* > 0 represents the activating strength of each activator and is obtained by training on the data. *θ*, also obtained by training, is the amount of activation in the absence of activator. *R* is the maximum synthesis rate, and we train on the target

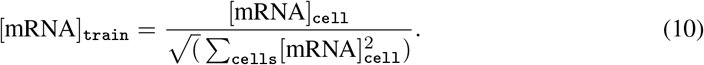

## 3 Implementation, Training, and Results

We implemented and trained this model in Keras with a TensorFlow back end ([10, 1]).The resulting architecture is shown in Figure 3. The resulting model is deemed by Keras to have 223 Layers and 52 unknown parameters. As this does not use any of the conventional architecture ([25, 19]), we frequently use the `Lambda` functions in Keras to represent some of the activation functions and PWM convolutions. Algorithm 1 was implemented as a special layer in Keras with two `rnn` functions.

**Figure 3:**
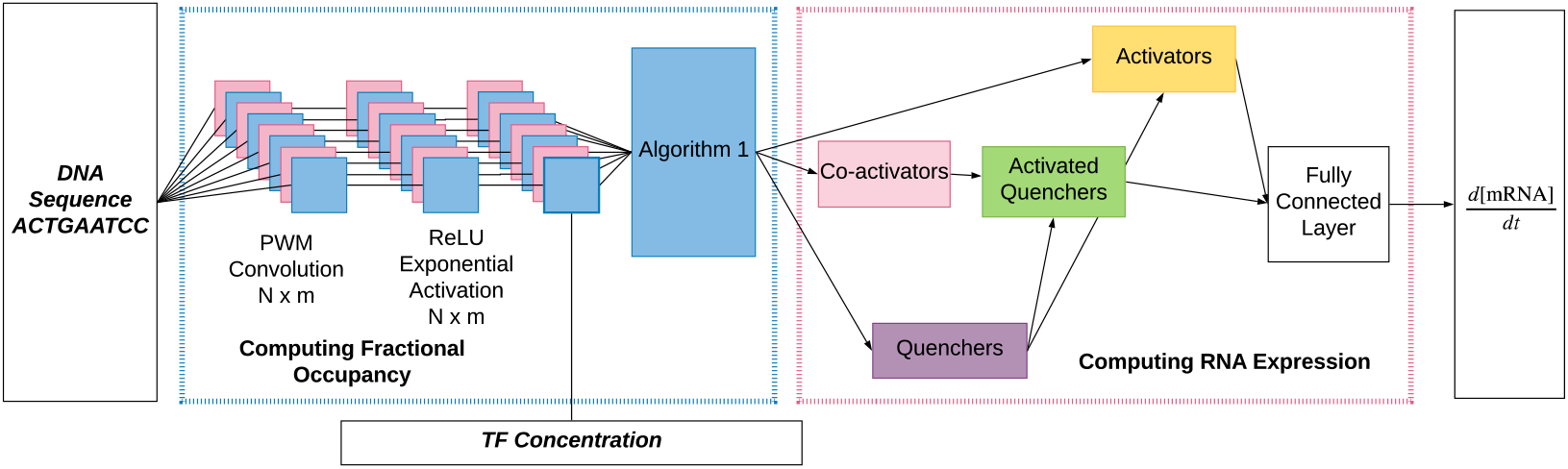
This is a graphical representation of the DNN. (Left) Chemical calculations. The computation can be seen as the DNA going through a convolution and then passing through a `ReLU` and then the the (exp(·)) activation function with bias. (Right) The graphical representation of interactions between bound TFs. Coactivators activate quenchers, quenchers quench activation, and activators combined together in a fully connected layer produce mRNA.

The training data and PWMs was as previously described [23], although training data was limited to the fusion constructs M32, M3_2, M23, and M2_3. The model was trained using a single `Intel Core i7-8700K CPU`. The training data contains 232 observations. The model was trained with 500 epochs using Adam ([24]) with Nesterov momentum as implemented in Keras. Training took approximately 4 hours. This compares with several days of serial simulated annealing before Algorithm 1 was devised [23], and is about equivalent to the time taken by code using Algorithm 1 running in parallel with the loss function for each construct computed on a separate core. However, optimization by simulated annealing requires several million evaluations of the loss function. while the implementation use here required about 10000. The earlier work used about 10000 lines of compiled C++ while this work uses less than 1000 lines of Python. However, the execution times indicate that there is considerable scope for improvement of the TensorFlow back end for this type of problem. Moreover, we see evidence of numerical issues in the Tensorflow back end which constrained us to make binding sites for all TFs the same size.

The results of the training are shown in Figure 2. We selected this training set, a pair of enhancers which produce greatly altered gene expression when fused, because they are known to produce a constrained model that is highly predictive [23]. It is important to note that this model tends not to over-fit the data since the number of parameters is only 52 compared to the 232 nuclei we used as the training set.

In a practical setting, the model should generate biologically accurate relative mRNA concentration at the right A-P position in % E.L. given a sequence not in the training set. We tested the predictive power of our DNN by confronting it with set of enhancer sequences which the model has not previously seen. In Figure 4, we show examples of four types of predictions, each of which consists of 58 nuclei and hence 58 separate predictions. We used the following enhancers for predictions. *run_str3_7* is the enhancer of the *runt* gene of *D.melanogaster* that drives *runt* stripes 3 and 7, each of which is about 2 % E.L. anterior of the corresponding eve stripes. Biologically, the prediction is reasonably accurate except for the low level of expression in the anterior. *eve_S2E(yak)* is the stripe 2 enhancer of *Drosophila yakuba,* expressed in *D. melanogaster* embryos [29, 30]. *D. yakuba* is a species of *Drosophila* that is closely related to *D. melanogaster* but has altered sets of binding sites. The position of Stripe 2 in this case is accurate but the level is low. *eve_S2E(cyn)* and *eve_S37E(put)* are respectively the enhancers of *eve* stripe 2 from *Sepsis cynipsea* and the 3/7 enhancer from *Themira putris,* both of the Sepsidae family, driving expression in *D. melanogaster* embryos [15]. These predictions are quite accurate including the fact that they are expressed slightly posterior to those of the *D. melanogaster eve* gene. These enhancers have a DNA sequence completely diverged from those of *D. melanogaster* [15, 14]. These predictions demonstrate the generalization capabilities of the DNN implementation of the model.

**Figure 4:**
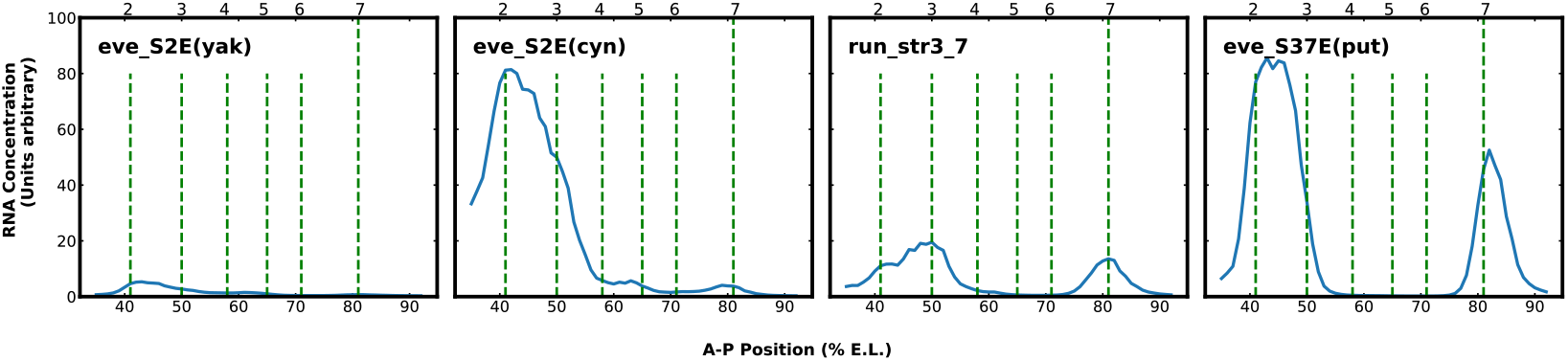
The figure shows four examples of predictions driven by enhancers not used for training. The location of eve stripes 2 through 7 are shown by vertical dashed lines. The vertical axis shows predicted mRNA concentration and horizontal axis shows A-P position in % E.L. The enhancers shown are described in the text.

## 4 Discussion

We discuss the the implication of the results presented here to both the biology and Deep Learning communities in turn. With respect to Deep Learning, our model constitutes an example of a fully interpretable DNN that is not merely biologically plausible but biologically validated [23]. It is our hope that this example will provide insights into the interpretabilities of DNNs in general, a problem that has received wide attention in the community [12, 8, 7, 26, 50, 31, 51].

More generally, neurobiology, together with the physics of spin-glasses, provided the initial inspiration for the mathematical structure of neural nets in general, and DNNs in particular were inspired by the structure of visual processing in the cerebral cortex. Perceptron based DNNs have an additive structure that arose from the additivity of voltages in neurons and spin glasses [33, 18]. In contrast, the mathematical structure of thermodynamic transcription models comes from multiplicative terms arising from the law of mass action, while the layered structure comes from the complex set of regulatory mechanisms that act in metazoan transcription. Perhaps the structure of the equations used here will suggest new applications and architectures for DNNs.

With respect to biology, the proof of concept presented here is much simpler to code than the original model, amounting to about 600 lines of Keras compared to about 10,000 lines of C++. Implementation in Keras also provides a numerical advantage by permitting the use of back propagation (BP) and stochastic gradient descent for optimization without the need to hand code partial derivatives. Although some issues connected to the speed of Keras implementation remain, these results suggest that models of this type could be scaled up to much larger datasets. Until recently such datasets, which must contain not only information about sequence but also data concerning the concentration of TFs, were scarce. Such datasets, on a genomic scale, have begun to be available for flies and mammals, including humans [3, 28, 46, 36, 49]. In these datasets, the functional role of TFs is typically unknown. In a study which used our model in mouse [6], all possible combinations of TF functional roles were considered, albeit with a considerable increase in computational expense. The data used in this study was not at a genomic scale, and an exhaustive search of TF functional roles at a genomic scale would be prohibitively expensive in computation. However, the introduction of reinforcement learning techniques may provide an avenue for surmounting this computational barrier. If the techniques presented here provide a way forward to precise characterization of gene regulation in metazoan organisms, it would be one further example of how investigation in the fruit fly *Drosophila*, with its special properties, can have global implications for biology.

## Acknowledgement

This work was supported by grant R01 OD10936 from the US NIH

